# Correlation of Alpha-1 Antitrypsin Levels and Exosome Associated Neutrophil Elastase Endothelial Injury in Subjects with SARS-CoV2 Infection

**DOI:** 10.1101/2022.07.07.499204

**Authors:** Jorge Lascano, Regina Oshins, Christina Eagan, Zerka Wadood, Xiao Qiang, Tammy Flagg, Yogesh Scindia, Borna Mehrad, Mark Brantly, Nazli Khodayari

## Abstract

**Background:** Severe acute respiratory syndrome caused by a novel coronavirus 2 (SARS-CoV-2) has infected more than 18 million people worldwide. The activation of endothelial cells is a hallmark of signs of SARS-CoV-2 infection that includes altered integrity of vessel barrier and endothelial inflammation.

**Objectives:** Pulmonary endothelial activation is suggested to be related to the profound neutrophil elastase (NE) activity, which is necessary for sterilization of phagocytosed bacterial pathogens. However, unopposed activity of NE increases alveolocapillary permeability and extracellular matrix degradation. The uncontrolled protease activity of NE during the inflammatory phase of lung diseases might be due to the resistance of exosome associated NE to inhibition by alpha-1 antitrypsin.

**Method:** 31 subjects with a diagnosis of SARS-CoV2 infection were recruited in the disease group and samples from 30 voluntaries matched for age and sex were also collected for control.

**Results:** We measured the plasma levels of exosome-associated NE in SARS-CoV-2 patients which, was positively correlated with the endothelial damage in those patients. Notably, we also found strong correlation with plasma levels of alpha-1 antitrypsin and exosome-associated NE in SARS-CoV-2 patients. Using macrovascular endothelial cells, we also observed that purified NE activity is inhibited by purified alpha-1 antitrypsin while, NE associated with exosomes are resistant to inhibition and show less sensitivity to alpha-1 antitrypsin inhibitory activity, in vitro.

**Conclusions:** Our results point out the role of exosome-associated NE in exacerbation of endothelial injury in SARS-CoV-2 infection. We have demonstrated that exosome-associated NE could be served as a new potential therapeutic target of severe systemic manifestations of SARS-CoV-2 infection.

## Introduction

Acute Respiratory Distress Syndrome (ARDS) is a life-threatening disease with an increasing incidence rate and approximately 30-40% rate of mortality [1]. Of hospitalized coronavirus 2019 (SARS-CoV2) patients, 33% develop some degree of ARDS [2], showing pulmonary inflammation, thick mucus airway secretions, extensive lung damage caused by elevated levels of pro-inflammatory cytokines secreted by activated immune cells, and thrombosis [3]. Excessive lung infiltration of activated neutrophils and release of neutrophil derived proteases are also hallmarks of ARDS [4]. Despite the role of neutrophils in repair of the inflamed tissues, neutropenia often does not result in improved recovery in ARDS patients [5].

Neutrophil Elastase (NE), a major protease released by neutrophils, involved in the inflammatory response, is released from activated neutrophils stimulated by cytokines and chemo-attractants at inflammatory sites. NE, a destructive elastase enzyme, modulates inflammation and tissue remodeling by proteolytic modification of cytokines and degradation of the extracellular matrix [6, 7]. It has been shown that a large part of released NE remains bound to the external surface of the plasma membrane where, it cleaves biologically relevant substrates [8]. Studies have shown that plasma levels of NE correlate with severity of lung injury in both animal models and human ARDS patients [9, 10].

Over 90% of free NE in an inflamed lung is rapidly inactivated by serine protease inhibitors such as alpha-1 antitrypsin (AAT), the most abundant serine protease inhibitor in the plasma [11]. Although several studies have demonstrated the levels of NE-AAT complexes as an indicator of NE activity, it does not systematically correlate with diseases severity [12]. AAT, a 52kDa glycoprotein synthesized primarily by hepatocytes [13], is an acute-phase protein, playing a critical role in limiting host-tissue injury mediated by proteases, particularly NE [14]. However, it has been shown that membrane-bound NE is resistant to inhibition by physiologic inhibitors, including AAT [8, 15]. Recently, a novel pathogenic entity of membrane-bound NE has been described wherein, activated neutrophils release NE associated with exosomes during degranulation which, are oriented in a configuration resistant to AAT [16].

Exosomes are a heterogeneous population of extracellular vesicles with an average diameter of around 150 nanometers which, have different biophysical functions in both physiological and pathological conditions [17]. Exosomes contain specific surface protein markers such as CD9, CD81, and CD63 [18]. Recently, studies have demonstrated that exosomes secreted from different cells transfer nucleic acids, proteins, or lipids to target cells, inducing changes in the recipient’s functions and phenotypes [17]. The effects of exosomes on target cells differ depending on their cargo, and such functional heterogeneity can result in different responses in different target cell types [19]. Given the biological properties of exosomes, the idea that circulating exosomes derived from inflamed tissue can modulate endothelial functions in response to inflammation is an active area of research [20].

During inflammation high concentrations of exosome-associated NE are likely to be present in inflamed tissues which, may not be efficiently inhibited by AAT, leading to NE-mediated tissue injury. NE-rich exosomes are thought to be secreted by activated neutrophils containing neutrophil-specific surface marker, CD66 [21]. Neutrophil-derived NE is suggested to play a pivotal role in the pathogenesis of ARDS mediated endothelial injury [22]. The pulmonary microvascular endothelium has unique role in homeostasis being frequently exposed to bloodborne pathogens, toxins, and endogenous inflammatory mediators. Under basal conditions, the quiescent and anti-inflammatory phenotype of endothelial cells is essential to lung homeostasis. Disruption of homeostatic endothelial functions can disturb barrier function, resulting in disrupted permeability that may lead to aberrant coagulation [23], which may predispose SARS-CoV-2 patients to organ failure in response to vascular injury.

The aim of the present study was to determine the abundance of exosome-associated NE in the plasma of SARS-CoV2 infected patients compared to healthy controls. We also investigated whether exosome-associated NE might bypass AAT and contribute to the pathophysiology of SARS-CoV2-mediated ARDS. In this study, we examined the plasma levels of free and exosome-associated NE, laboratory markers of endothelial viability such as CRP and LDH [24][25], and endothelial integrity markers such as PECAM-1 from SARS-CoV2 infected patients to investigate the involvement of exosome-associated NE in worse outcomes in those with severe illness.

## Materials and methods

### Study Population

Prospective subject enrollment occurred at UF Health hospital. Subjects were recruited from the different hospital floors including the medical intensive care unit as well as regular hospitalization floors. All patients were admitted with a diagnosis of SARS-CoV2 infection. Confirmation of the diagnosis was done by PCR. 31 subjects were recruited in the disease group and samples from 30 voluntaries matched for age and sex were also collected for control. All subjects or appointed health care surrogate provided written informed consent. The only exclusion criteria was inability to obtain consent. All patients gave informed, written consent prior to enrolling. The study was approved by the University of Florida institutional IRB and is listed at clincaltrials.gov (IRB # 202000779).

### Blood Sample Collection

Blood samples were collected from peripheral venipuncture with a butterfly, in two 3.2% buffered Na citrate tubes (Becton Dickinson). Samples were taken immediately to the alpha-1 antitrypsin deficiency laboratory at the University of Florida for process and storage. Blood was centrifuged at 2000 x g for 10 minutes at room temperature after which the plasma was collected, aliquoted, and stored at -80° C.

### Isolation of Plasma Exosomes

Exosomes were isolated from 300 µl of plasma from 30 healthy individuals and 30 subjects with SARS-CoV2 infection using “Total Exosome Precipitation Reagent” from Invitrogen (Carlsbad, CA). Briefly, 300 µl of clarified plasma was mixed with 150 µl of 1X PBS. Then, 90 µl of the Exosome Precipitation Reagent was added to the sample and mixed by vortex. The samples were incubated at room temperature for 10 minutes followed by centrifugation at 10,000 × g for 5 minutes at room temperature. The pellet containing the exosomes was resuspend in 1X PBS and conserved at − 80 °C for later use [17].

### Characterization of Plasma Exosomes

The size distribution and concentration of the isolated exosomes was analyzed by the NanoSight NS300 system (NanoSight and Malvern, United Kingdom). Briefly, purified exosomes were homogenized followed by dilution of 1:100 in sterile PBS. Each sample analysis was conducted for 60 seconds. Data was analyzed by NanoSight NTA 2.3 Analytical Software with the detection threshold optimized for each sample and screen gain at 10 to track as many particles as possible with minimal background. A blank 0.2 μm-filtered 1x PBS sample was also run as a negative control. Five analyses were done per sample [17].

### Determination of Free and Exosome-Associated Neutrophil Elastase

Extracted exosomes were lysed with RIPA buffer and treated with 1 mM PMSF for 30 minutes at room temperature. The exosome-free plasma supernatant from the extraction was treated with PMSF only. For the assay, sheep anti-human NE antibody (US Biological Life Sciences, Salem, MA) was nonspecifically adsorbed to microtiter wells overnight at 4°C before samples were applied and incubated at 37°C for 2 hours. Plates were washed and samples incubated with rabbit anti-human NE (Athens Research and Technology, Athens, GA) for 1 hour at 37°C. Plates were washed and incubated with HRP-conjugated goat anti-rabbit IgG (MilliporeSigma, St. Louis, Missouri) for 1 hour at 37°C. Samples were developed using TMB substrate and absorbance read using a SpectraMax M3 spectrophotometer at 450 nm.

### Determination of Vascular Endothelial Inflammatory Markers

Plasma from 30 healthy individuals and 31 SARS-CoV2 infected patients was analyzed using the Milliplex Human Cardiovascular Disease Magnetic Bead (MilliporeSigma, St. Louis, Missouri) according to manufacturer’s instructions. Briefly, samples were combined with serum matrix buffer and magnetic beads labeled for sE-Selectin, PECAM-1, Pentraxin-3, Tissue Factor (TF), and Thrombomodulin incubated overnight on a shaker at 4°C. The beads were washed and incubated with detection antibody for 1 hour followed by streptavidin-phycoerythrin for 30 minutes at room temperature. The beads then were washed, and mean fluorescence intensity was measured using a Luminex instrument (Invitrogen, Waltham, MA).

### Cell Culture and Treatment Protocols

A human HL-60 cell line (ATCC) was cultured in RPMI containing 10% FBS, 1% Primocin, and Insulin-Transferrin-Selenium (ITS-G, 1X, ThermoFisher, Waltham, MA). For differentiation into neutrophil-like cells, the media was supplemented with 1.3% DMSO and the cells grown for 5 days. Cells were then treated with either 1µM fMLP (N-formyl-L-methionyl-L-leucyl-phenylalanine, Sigma) or DMSO overnight in media containing 10% exosome-free FBS. Exosomes were collected from the media by ultracentrifugation. Human lung microvascular endothelial cells (MVEC) from healthy donors [26], a gift from Dr. Andrew J. Bryant, were cultured on collagen in Vascular Cell Basal Medium supplemented with the VEGF Endothelial Cell Growth Kit (ATCC, Manassas, VA). Cells were treated with either 1×10^7^ per well of quiescent exosome from HL-60 cells, activated (fMLP treated) exosomes from HL-60 cells [17], 34 nM of purified neutrophil elastase [27, 28] (Athens Research and Technology. Athens, GA), 34 nM of neutrophil elastase and purified alpha-1 antitrypsin (Grifols USA, Los Angeles, CA), or activated HL-60 exosomes and alpha-1 antitrypsin (34 nM) overnight before RNA was collected using Trizol.

### Immunoblotting

Exosome lysates were separated by 4-20% SDS-PAGE and transferred to a nitrocellulose membrane. Membranes were blotted with specific antibodies to CD66, CD63, and CD81 (ProteinTech, Rosemont, IL) and neutrophil elastase (R&D systems, Minneapolis, MN).

### Gene Expression Measured by Real-Time PCR (RT-PCR)

RT-PCR was conducted using Applied Biosystems TaqMan commercial primers on an ABI Prism 7500 fast detection system using standard protocol; 18S mRNA was used as an internal reference. Quantification of relative gene expression was performed using the comparative threshold cycle (CT) method [29].

### Statistical Analysis

All results from the study are expressed as mean ± S.D. Statistical analyses were performed using Prism 8 by Student’s t-test or Mann-Whitney U test. Values of *p* < 0.05 were considered statistically significant.

## Results

### Baseline characteristics of the study population

The demographic and clinical characteristics of the COVID-19 patients are shown in Table 1. The mean (+/-SD) age of the overall SARS-CoV2 cohort was 63.8 years and 35% were men. The most common symptoms on admission to the hospital were fever, dyspnea and cough. Comorbidities were common in the study population. Over 23% of the total cohort had pre-existing lung disease, 68% had hypertension, 16% had documented coronary artery disease, 48% had diabetes mellitus and 32% of patients were smokers. ARDS severity was defined by Pao2/Fio2 ratio. 22.5% of patients did not have ARDS, 35.4% had mild ARDS, 29% had severe ARDS and 13% had sever ARDS (Fig 1A). Circulating levels of AAT (Fig 1B) and CRP (Fig 1C) were elevated (42 ±12.96, and 178 ±120.9 respectively) in patients with a diagnosis of COVID-19 compared to Healthy controls.

**Table 1.** Baseline characteristics of the study population.

| Patients Demographics n (31) |  | Controls Demographics n (29) |
| --- | --- | --- |
| **Age mean(+/-SD)** | 63.8 (11.9) | 45.6 (15.5) |
| **Gender** |  |  |
| Male | 35% | 52% |
| Female | 65% | 48% |
| **Race** |  |  |
| Caucasian | 32% | 48% |
| African American | 29% | 4% |
| Other/Unknown | 39% | 48% |
| **ARDS by (Pao2/Flo2)** |  | N/A |
| No ARDS | 22.5% |  |
| Mild ARDS | 35.4 |  |
| Moderate ARDS | 29% |  |
| Severe ARDS | 13% |  |
| **BMI mean (+/-SD)** | 37.5 (9.1) | 24.8 (3.4) |
| **ICU/ Non-ICU location** | 85% / 15% | N/A |
| **APACHE mean(+/-SD)** | 11.6 (4.1)) | N/A |
| **SOFA mean(+/-SD)** | 3.7 (1.8) | N/A |
| **Comorbidities** |  |  |
| Hypertension | 68% | N/A |
| Diabetes | 48% | N/A |
| CAD | 16% | N/A |
| Obesity | 74% | N/A |
| Chronic lung disease | 23% | N/A |
| Tobacco use | 32% | N/A |

**Figure 1.**
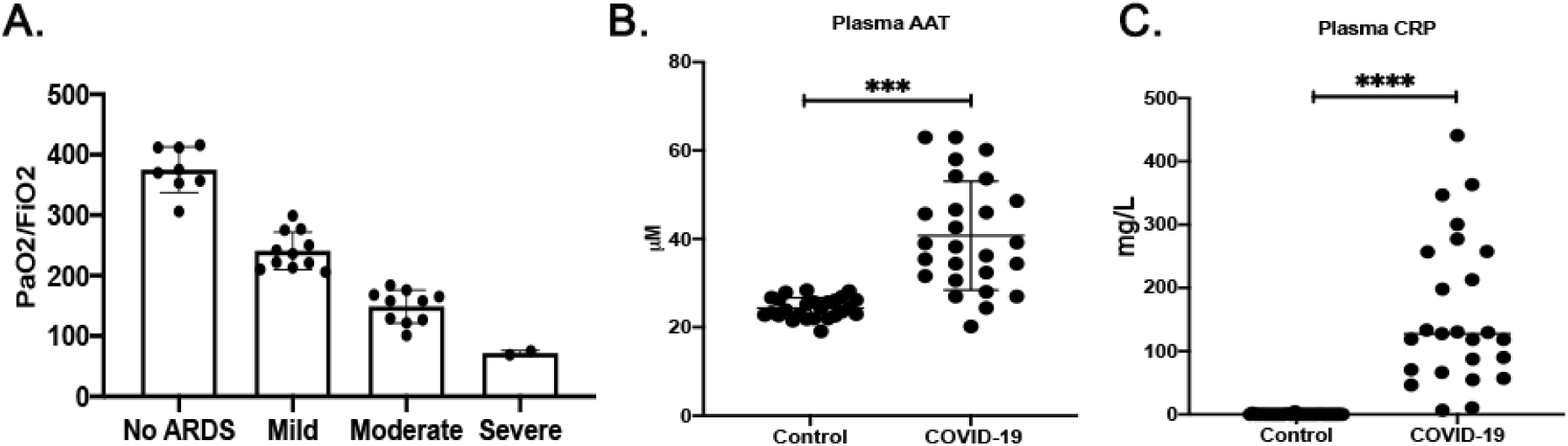
Baseline inflammation in COVID-19 patients compared to Healthy controls. (A) The levels of PaO2/FiO2 and ARDS scoring in COVID-19 patients. (B) circulating AAT and (C) CRP in COVID-19 patients compared to Healthy controls, ***p<0.0005, #x002A;***p<0.0001.

### Correlations among plasma profiles in the SARS-CoV2 patients

We used PaO2/FiO2 ratio to assess lung function in SARS-CoV2 patients as well as the plasma levels of lactate dehydrogenase (LDH), CRP, and AAT as prognostic markers reflecting acute phase reactions. As it was expected [30, 31], there was highly significant correlation between plasma levels of CRP and AAT from COVID-19 patients (Fig 2A). Lower levels PaO2/FiO2 were negatively correlated with elevated levels of LDH which was reported previously as an indication of mortality rate [32, 33] (Fig 2B). Consistent with previous reports, we also observed Higher levels of LDH in non-survivors compared with survivors (411.3±160.2, and 345.1±131.3, respectively) (Fig. 2C), which were also modestly correlated with higher plasma levels of CRP and AAT in all patients (Fig 2D and E) and severity of ARDS based on PaO2/FiO2 ratio values (Fig 2F). Furthermore, we were not able to see significant differences in the plasma levels of AAT and CRP in non-survivors compared with survivors (S1 Fig 1A and 1B), and significant correlations between PaO2/FiO2 and CRP or AAT in our study subjects (S1 Fig 1C and 1D).

**Figure 2.**
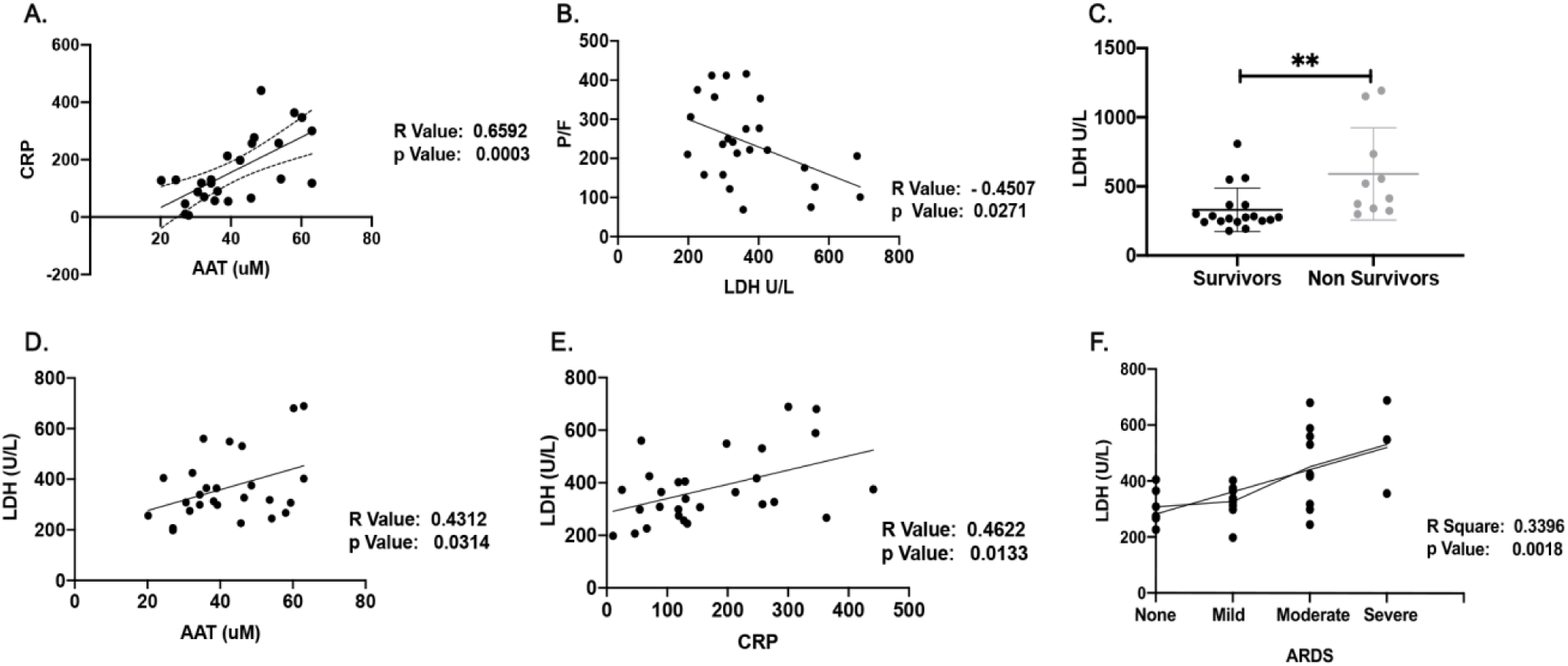
Correlations among plasma profiles in the SARS-CoV2 patients. (A) The correlation between plasma levels of CRP and AAT from COVID-19 patients. (B) Negative correlation between PaO2/FiO2 levels and plasma LDH levels. (C) Plasma LDH levels as indication of mortality rate. (D and E) The correlation of LDH levels and plasma levels of CRP and AAT. (E) Severity of ARDS based on PaO2/FiO2 ratio values, **p<0.005.

### Plasma levels of exosome associated NE increase with the clinical progress of SARS-CoV2-mediated ARDS

Given that the relationship observed between plasma levels of CRP and AAT in SARS-CoV2 patients was consistent with disease severity, we searched for evidence of an altered inflammatory mediator in these individuals. We examined the plasma circulating exosomes from healthy normal individuals compared to SARS-CoV2 patients for surface bound exosomal NE. The result showed that SARS-CoV2 patients had significantly higher plasma levels of exosome associated NE (2587±2372) compared with healthy controls (777.3±1195) (Fig 3A). The association between plasma levels of exosome associated NE and plasma exosome associated levels of CD66 as a marker of neutrophil derived exosomes, was assessed by western blot analysis (Fig 3B). Furthermore, the plasma levels of exosome associated NE in SARS-CoV2 patients was significantly correlated with plasma levels of free NE (Fig 3C). There were also moderate correlations between the plasma levels of exosomal NE and CRP and AAT levels (Fig 3D and E).

**Figure 3.**
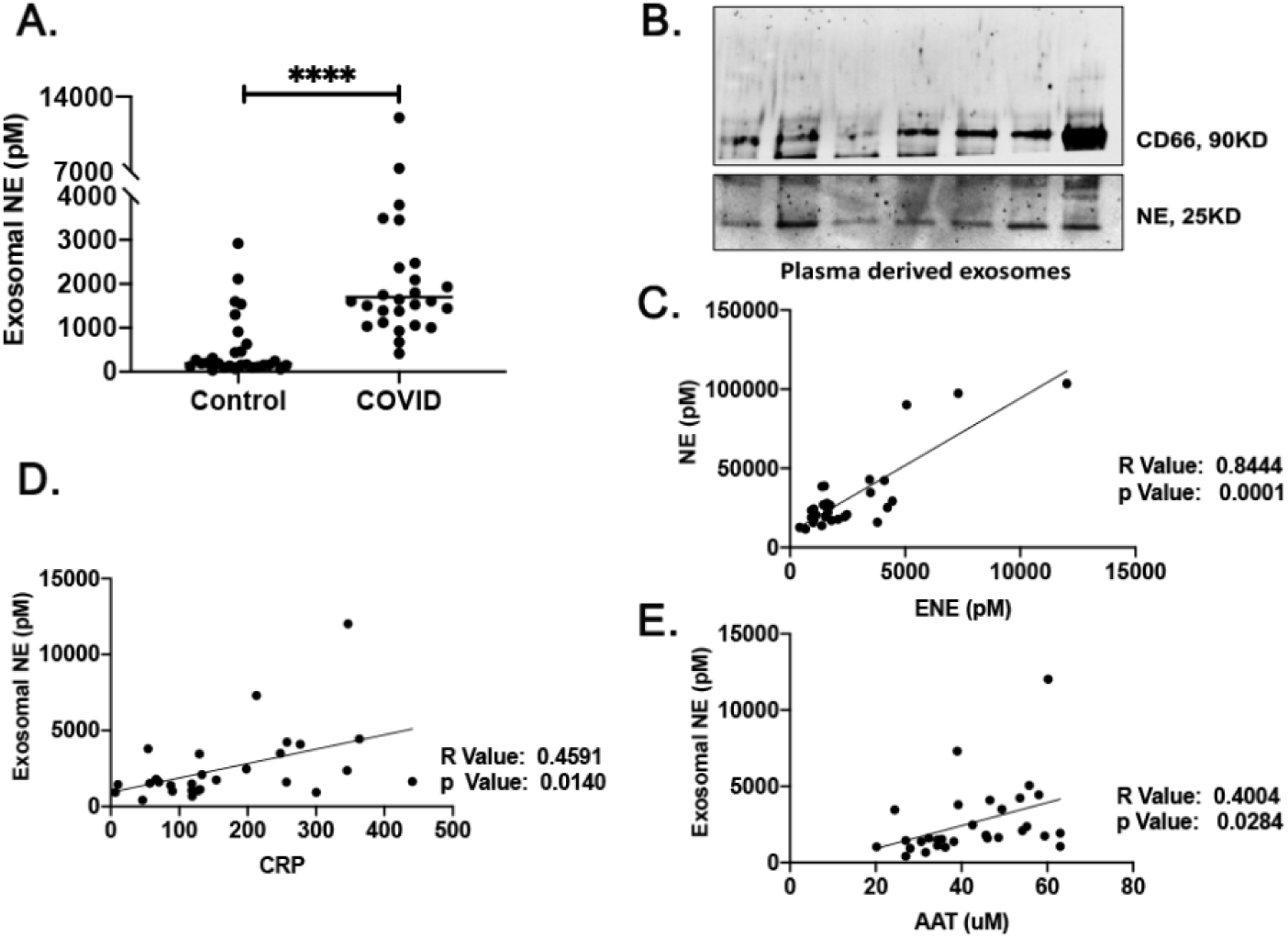
Plasma levels of exosome associated neutrophil elastase (NE) in SARS-CoV2 patients. (A) Higher plasma levels of exosome associated NE in SARS-CoV2 patients compared with healthy controls. (B) The association between exosome associated NE and plasma exosome associated levels of CD66 in SARS-CoV2 patients assessed by western blot analysis. (C) The correlation of the plasma levels of exosome associated NE (ENE) in SARS-CoV2 patients with levels of free NE. (D) The correlation of the plasma levels of exosome associated NE in SARS-CoV2 patients with levels of CRP, (D) and AAT, ****p<0.0001.

### Correlations among plasma exosome associated NE and endothelial injury in the SARS-CoV2-mediated ARDS patients

We next sought to investigate the relation between plasma exosome associated NE and endothelial injury in SARS-CoV2 patients. The result showed that in SARS-CoV2 patients higher plasma levels of exosome associated NE correlated with the plasma levels of LDH (Fig 4A). Owing to the alterations in inflammatory markers observed in SARS-CoV2 patients, we also investigated changes in soluble PECAM-1 plasma levels. SARS-CoV2 patients had a two-to three-fold increase in mean soluble PECAM-1 plasma levels (64.42±42.84) compared with healthy control (16.27±10.54) individuals (P <0.0001) (Fig 4B). Results also indicated that plasma soluble PECAM-1 values were correlated with the levels of plasma exosome associated NE (Fig 4C) and negatively correlated with the PaO2/FiO2 ratio (Fig 4D). There was also a moderate correlation between the plasma soluble PECAM-1 values and plasma levels of AAT in COVID-19 patients (Fig 4E). In addition to PECAM-1 plasma levels, we have also observed an increase in the plasma levels of Tissue Factor (TF) in SARS-CoV2 patients compared to controls (5.258±2.75, and 2.347±1.943 respectively) (Fig 4F) which was moderately correlated with plasma levels of exosome associated NE (Fig 4G).

**Figure 4.**
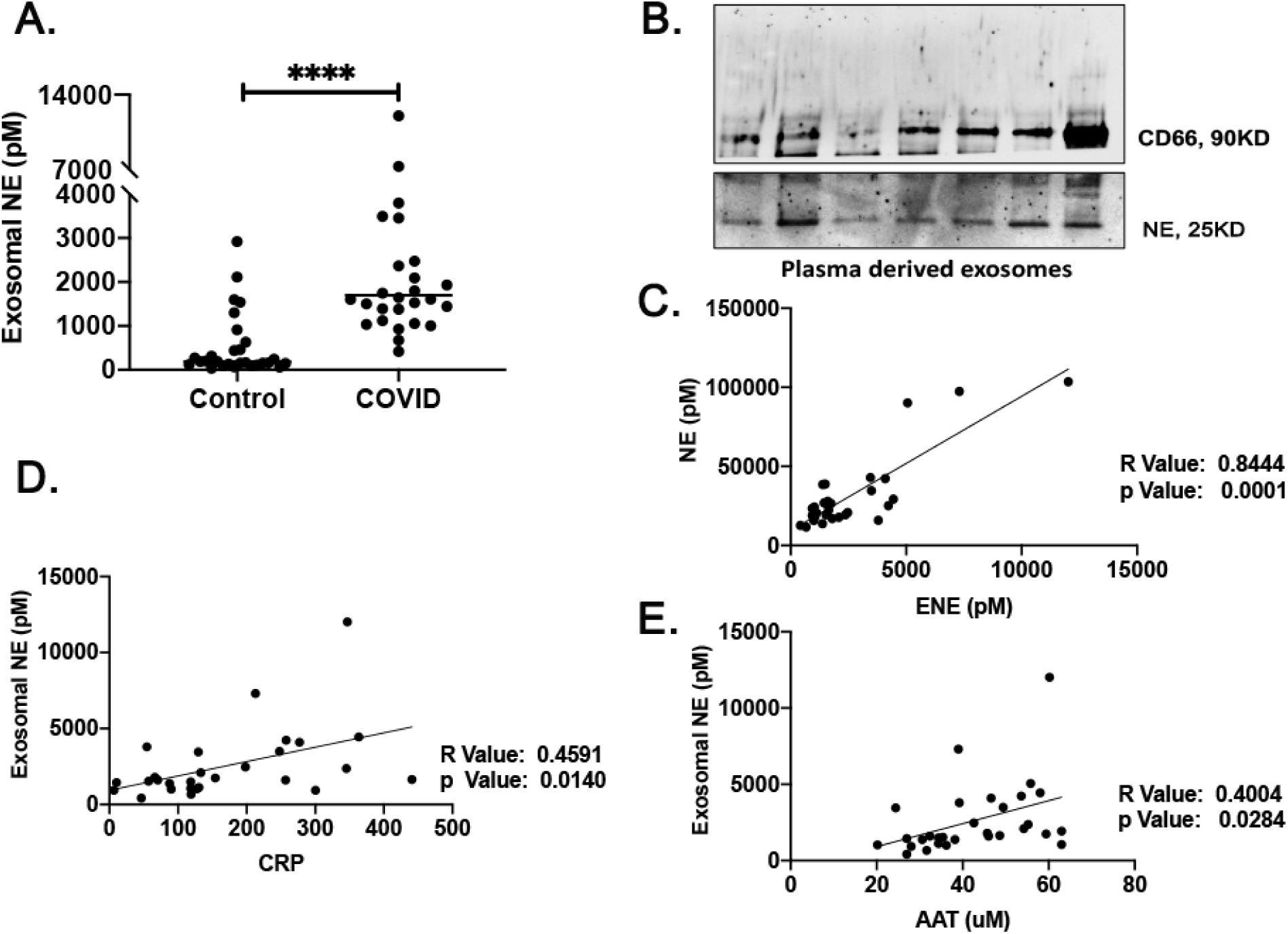
Correlations among plasma exosome associated neutrophil elastase (NE) and endothelial injury in the SARS-CoV2-mediated ARDS patients. (A) The correlation of plasma levels of exosome associated NE levels of LDH in SARS-CoV2 patients. (B) Soluble PECAM-1 plasma levels of SARS-CoV2 patients compared with healthy controls. (C) The correlation of plasma soluble PECAM-1 values with plasma exosome associated NE. (D) Negative correlation of PaO2/FiO2 ratio with soluble PECAM-1 values and (E) correlation between the plasma soluble PECAM-1 values and plasma levels of AAT in COVID-19 patients. (F) Plasma levels of Tissue Factor (TF) in SARS-CoV2 patients compared with healthy controls. (G) The correlation of plasma TF values with plasma exosome associated NE. ****p<0.0001.

### Exosomes from activated neutrophil-like cells carry surface NE

We examined exosomes released by neutrophil-like cells activated with the bacterial formylated peptide (a canonical neutrophil stimulant) formyl-methionine-leucine-phenylalanine (fMLP), referred to here as “activated exosomes,” and those released constitutively from DMSO control-treated neutrophils (“quiescent exosomes”). The size and concentration of the exosomes was assessed using NTA analysis [34] (Fig 5A), which showed quiescent and activated exosomes were similar in size (∼150 nm) and quantity released (Fig 5B and C). Both exosome populations expressed CD66, a neutrophil-associated protein, but general exosome marker CD81 was more abundant in activated exosomes (Fig 5D). Using ELISA, we found NE was present on the exosome surface and confirmed that activated exosomes had considerably higher quantities of surface NE compared to quiescent exosomes (Fig 5E).

**Figure 5.**
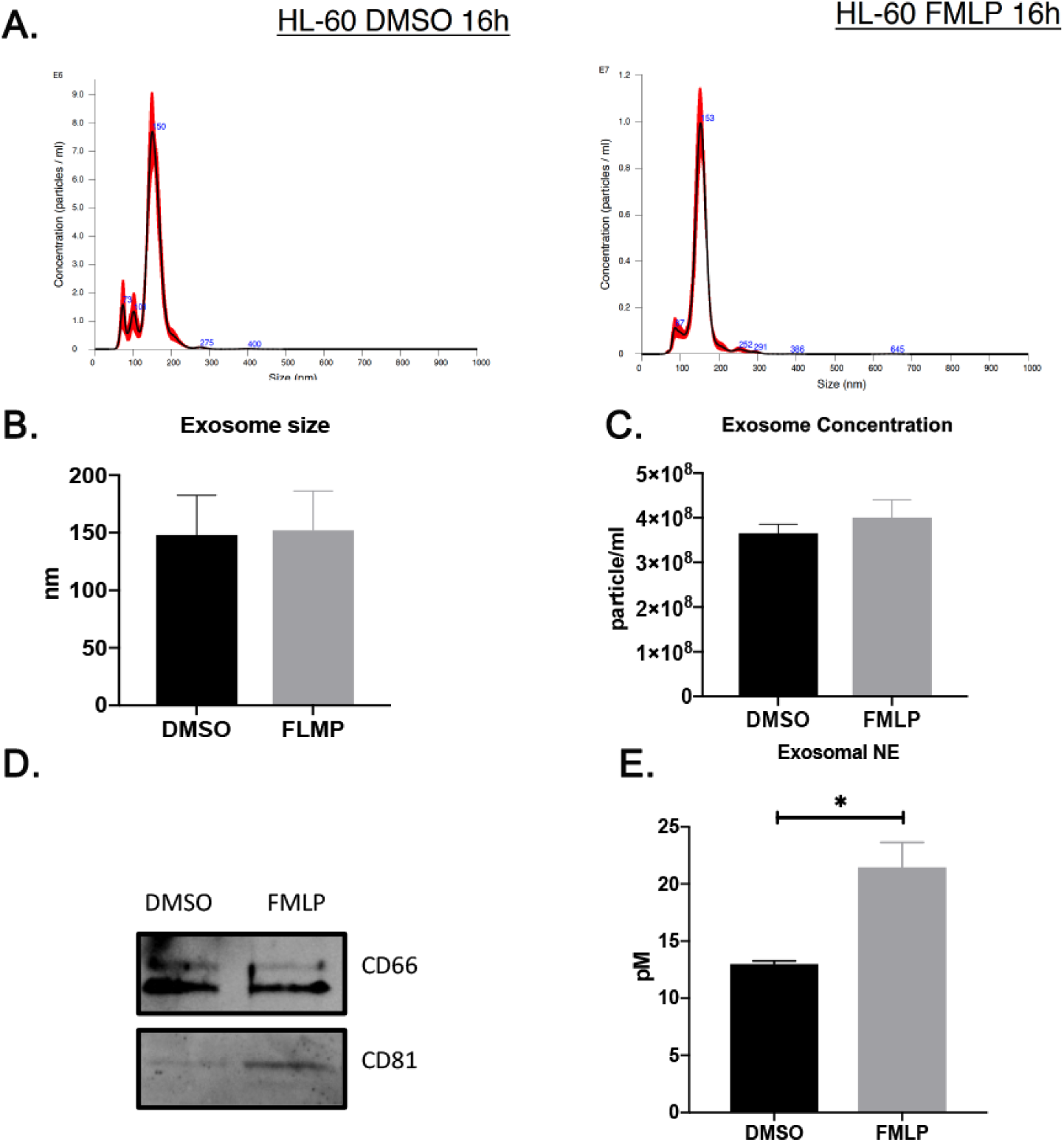
Exosomes from activated neutrophil-like cells carry surface neutrophil elastase (NE) in SARS-CoV2 patients. (A) The size and concentration of the exosome from activated quiescent exosomes, assessed using nano tracker. (B) Similar size and (C) concentration of the quiescent and activated exosomes. (D) Exosomal markers of CD66 and CD81 assessed by western blot analysis. (D) The levels of NE in the exosomal fractions assessed by ELISA, *p<0.05.

### Unchecked activity of exosome associated NE activates lung endothelial cells in vitro

We next sought to test whether the activity of exosome associated NE applied to lung microvascular endothelial cells (MVECs), activating them in vitro. RNA expression levels of different pro-inflammatory cytokines in the MVECs were analyzed 8 hours post treatment with purified NE, quiescent exosomes, and activated exosomes with or without purified AAT using qPCR. Both purified NE and activated exosomes were able to increase the RNA expression levels of IL-6 and IL-8 in MVECs after 8 hours of incubation. Our results also showed that purified NE activity was largely inhibited by purified AAT. In contrast, NE associated with activated exosomes was resistant to inhibition by purified AAT and showed less sensitivity to AAT inhibitory activity (Fig 6).

**Figure 6.**
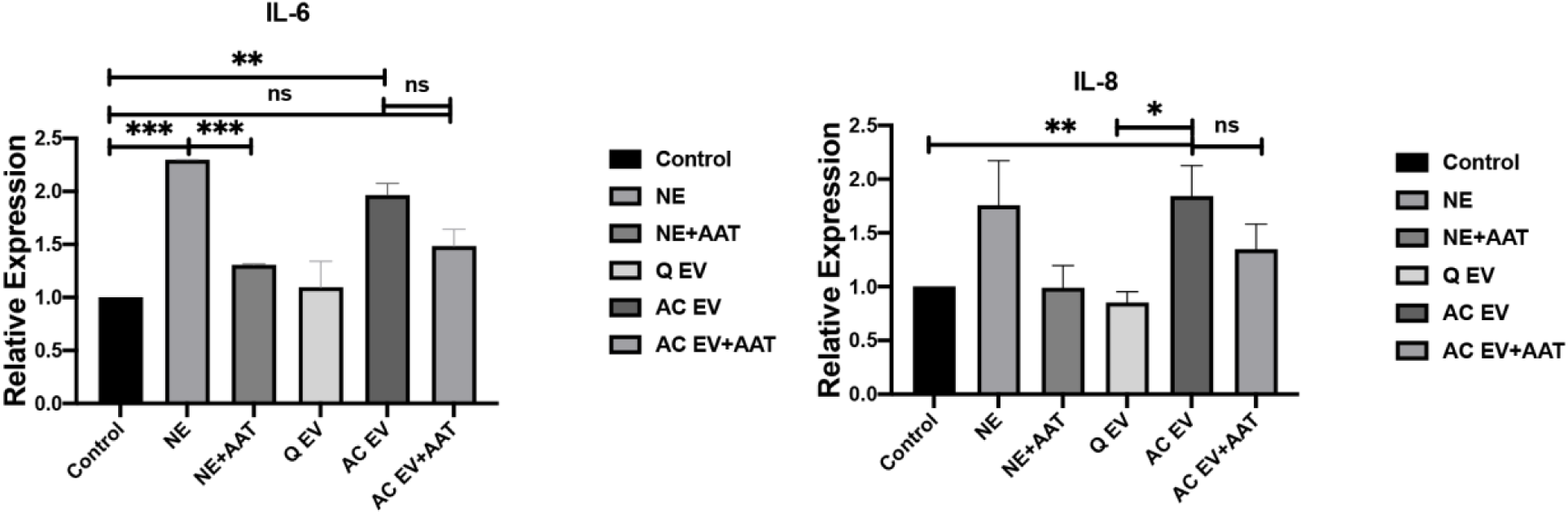
Unchecked activity of exosome associated neutrophil elastase (NE) activates lung endothelial cells in vitro. RNA expression levels of different pro-inflammatory cytokines in the MVECs analyzed 8 hours post treatment with purified NE, quiescent exosomes, and activated exosomes with or without purified AAT using qPCR, *p<0.05, **p<0.005, ***p<0.0005.

## Discussion

An imbalance between proteinases and their physiological inhibitors in the lung plays a pivotal role in the initiation and propagation of acute lung injury such as SARS-CoV2-mediated ARDS [11]. In this study we extend this observation into the peripheral circulation. Here, we demonstrate signs of COVID-19 mediated endothelial injury in patients with SARS-CoV2 infection and show the correlation of endothelial phenotype with increased plasma levels of exosome-associated NE. In addition, our results show that exosome-associated NE is not inhibited by AAT and is able to activate lung endothelial cells. Consistent with previous reports [35], we found the most unanticipated factor among patients with severe COVID-19 is a relatively blunted AAT protease inhibitory response.

Many experimental and clinical studies have indicated that enhanced activity of NE is associated with the pathogenesis of acute and chronic inflammatory diseases [36]. NE, a neutrophil-derived serine protease, plays a crucial role in tissue damage during chronic inflammatory course of the diseases such as ARDS [37]. NE activity is associated with severity of lung disease and surface-bound NE activity correlates with pulmonary function indices of airflow limitation and pulmonary hyperinflation [38]. It has been also shown that exosome-associated NE activity is not inhibited by endogenous antiproteases such as AAT and plays an important role in the pathogenesis of chronic neutrophilic lung diseases. The excessive activity of exosome-associated NE leads to damaged extracellular matrix and endothelial tissue. As a consequence, patients with such exosomes in circulation may be viewed as functionally AAT-deficient and may have exosome associated NE-driven inflammation in addition to local elevations of NE [16].

We have demonstrated that patients with COVID-19 displayed increased AAT plasma levels compared to healthy controls. This is correlated with elevations in plasma CRP, a marker of systemic inflammation [39], and LDH, a marker of cellular injury [40]. It has been shown that plasma AAT levels in the presence of inflammation are associated with serum CRP levels [39]. In line with previous reports [30], we have found that in COVID-19 patients, plasma LDH and CRP levels correlate with respiratory failure based on PaO2/FiO2 ratios. We have also shown that plasma levels of LDH may serve clinically as a prognostic survival factor in SARS-CoV2-ARDS patients as it significantly correlates with the severity of ARDS. LDH, an inflammatory marker involved in cell’s energy production, is present in almost all cells with highest levels in the heart, liver, and lung tissues [41] and has been recently shown to be highly correlated with NE in COVID-19 [42]. CRP, a general marker of mild systemic inflammation, is a hepatic protein regulated by cytokines IL-6 and IL-1 [43]. The most interesting findings of our study were elevated levels of plasma exosome-associated NE in COVID-19 patients compared to healthy controls and its correlation with elevated levels of AAT, CRP, and LDH concentrations in the plasma.

Exosomes from activated neutrophils express increased amounts of enzymatically active surface NE which is capable of unchecked proteolytic activity [16]. In inflammatory diseases, the excessive activity of neutrophil-derived exosomes leads to damaged extracellular matrix, resulting in alveolar loss. These exosomes may also contribute to pulmonary architectural distortion seen in lung diseases, including ARDS, which is associated with excessive neutropenia-mediated inflammation and elevation of NE levels [16]. Alternatively, endothelial cells could be directly affected by proteolytic activity of exosome associated-NE and contribute to endothelial injury [44].

A variety of endothelial cell surface adhesion molecules have been shown to also exist as soluble proteins which, are thought to originate by cleavage of the cell surface adhesion molecules. Platelet-endothelial cell adhesion molecule 1 (PECAM-1) is one such soluble adhesion molecule which is elevated in a variety of inflammatory diseases [45]. PECAM-1, highly expressed at endothelial intercellular junctions as well as in platelets and immune cells, has a regulatory role in leukocyte trafficking and cell junctional integrity maintenance [46]. Under normal conditions, levels of soluble PECAM-1 are very low, which is indicative of normal functioning. Higher circulating levels of PECAM-1, as occurring during inflammatory diseases such as ARDS, may serve as a diagnostic marker of endothelial dysfunction [47]. Little is known about the mechanisms involved in PECAM-1 expression and processing in endothelial cells. Current evidence suggests activated inflammatory cells undergo an active shedding of the extracellular domain of the PECAM-1 molecule during inflammation which could contribute to the loss of cell-to-cell adhesion [47]. It has been shown that MMP-9 (Matrix metalloproteinase-9) mediated cleavage of PECAM-1 leads to increased levels of soluble PECAM-1 in vivo [48], which indicates MMP-dependent mechanisms underlie the endothelial injury mediated by inflammation. In our study we show there is a correlation between increased plasma levels of PECAM-1 in SARS-CoV2 patients and the plasma levels of exosome-associate NE, a potent physiological activator of MMP-9 [49]. This supports the concept of exosome-associated NE involvement in the pathogenesis of increased endothelial permeability seen in SARS-CoV2-induced ARDS.

It has been previously observed that plasma TF levels may be the risk factor for thrombosis [50]. TF synthesis and release is stimulated by cell damage which can induce the proliferation and migration of endothelial cells, and it is involved in inflammation [51]. Consistent with previous reports [52], we also have observed elevated plasma levels of TF in our SARS-CoV2 study cohort compared to controls which was mildly correlated with plasma levels of exosome-associated NE. The endothelial cell’s damage mediated by proinflammatory cytokines [53] as well as exosome-associated NE may lead to direct contact between TF and circulation, which in theory may result in increased levels of TF in the plasma of these patients. Therefore, this issue certainly requires more study in the future.

Although substantial evidence supports the role of NE in the lung microvascular dysfunction in ARDS [54-57], plasma and lung interstitial fluid contain proteinase inhibitors, such as AAT, that normally protect tissue from the unregulated action of NE [58, 59]. Consistent with recent demonstration [16, 60], here we show the activity of exosome-associated NE is resistant to AAT inhibitory functions and leads to activation of endothelial cells in vitro. The results suggest that such a mechanism may contribute to the pulmonary architectural distortion seen in other lung diseases, that are associated with local elevations of NE and functionally AAT deficiency.

Our study has several limitations. First, we were only able to compare our findings to normal controls. Ideally, we would have liked to use non-covid ARDS/Pneumonia subjects also, but we did not have access to such patients during our recruitment time. Others including McElvaney et al [35] have shown that the inflammatory profile of subjects with COVID-19 infection admitted to the ICU is different than the profile of non-covid Pneumonia admitted to the ICU. Second our sample size is small leading to some degree of variability that is difficult to control.

In conclusion, this study contributes to the expanding knowledge of the pathophysiology of SARS-CoV2 mediated ARDS disease. We suggest that plasma exosomes released by activated neutrophils play a role in COVID-19 pathophysiology, contributing to the augmented pro-inflammatory response and endothelial dysfunction observed in COVID-19 patients. Our findings collectively suggest that the efficacy of protease inhibitors against the exosome-associated form of NE maybe an important consideration in therapeutics development against endothelial injury in lung diseases. Perhaps this knowledge around exosome associated NE will offer a novel way to approach therapies to treat this disease and other inflammatory related illnesses.

## Abbreviations

AAT: Alpha-1 Antitrypsin
ARDS: Acute Respiratory Distress Syndrome
CRP: C-Reactive Protein
fMLP: Formyl-methionine-leucine-phenylalanine
LDH: Lactate Dehydrogenase
MVEC: Lung Microvascular Endothelial Cells
NE: Neutrophil Elastase
PECAM-1: Platelet-endothelial cell adhesion molecule 1.

## Availability of data and materials

All data generated or analyzed during this study are included in this published article.

## Declarations of interest

The authors have no conflicts of interest to declare.

## Author contributions

J.L. – study concept and design, analysis and interpretation of data, drafting of manuscript, study supervision. R.O. – acquisition of data, technical support. C.E – acquisition of data. Z.W – acquisition of data. X.Q. – technical support. T.F. – technical support. Y.S. – technical support. B.M. – analysis and interpretation of data, critical revision of manuscript for important intellectual content. M.B. – study concept and design, analysis and interpretation of the data, critical revision of manuscript for important intellectual content, study supervision. N.K. – study concept and design, acquisition of data, analysis and interpretation of data, statistical analysis, drafting of manuscript, study supervision. All authors read and approved the final manuscript.

## Notes

### Competing Interest Statement

The authors have declared no competing interest.

