## Supplementary figures and images for "Correlation of Alpha-1 Antitrypsin Levels and Exosome Associated Neutrophil Elastase Endothelial Injury in Subjects with SARS-CoV2 Infection"

### Supplemental Figure1

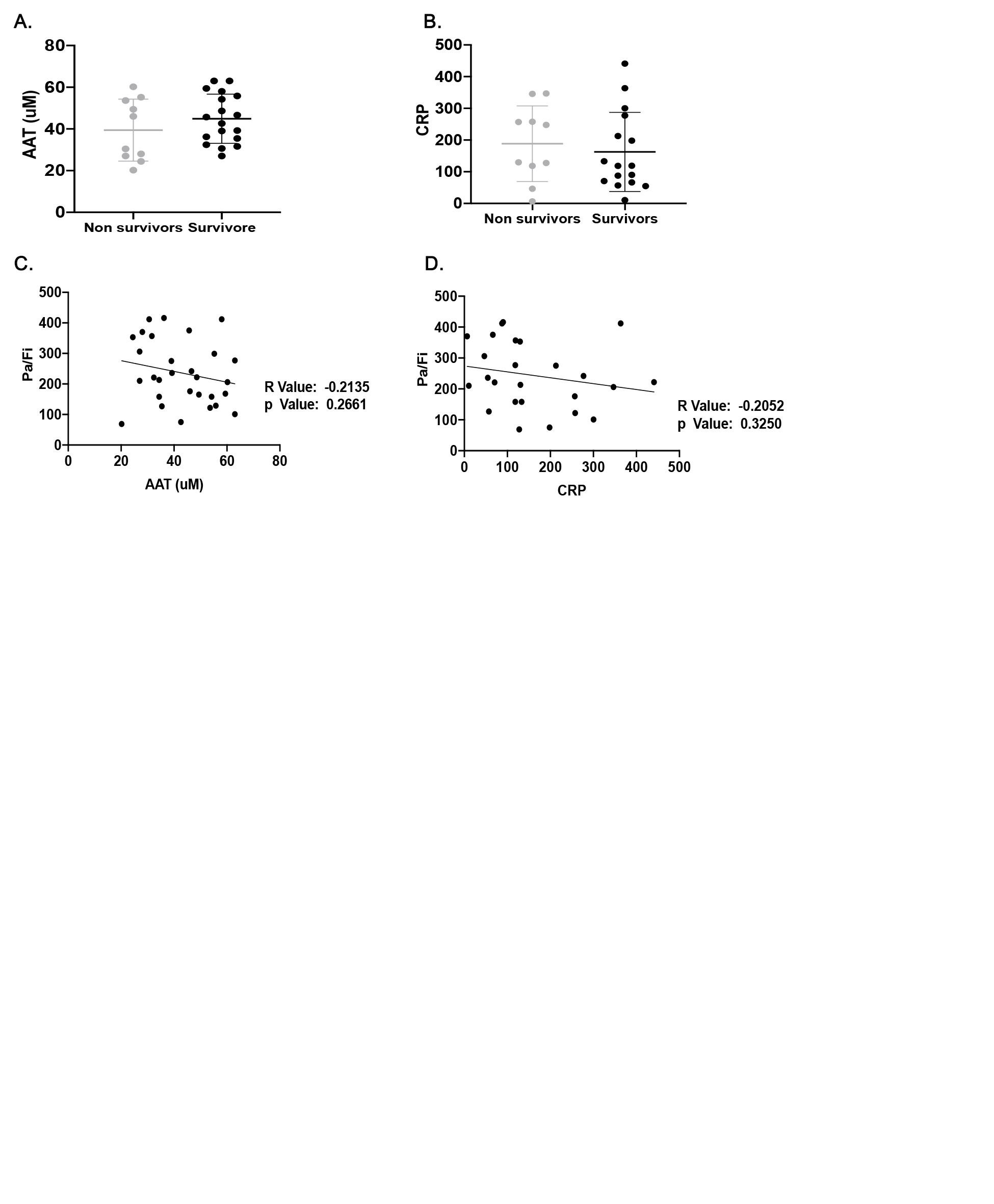
